# Cecelia: a multifunctional image analysis toolbox for decoding spatial cellular interactions and behaviour

**DOI:** 10.1101/2024.08.13.607845

**Authors:** Dominik Schienstock, Jyh Liang Hor, Sapna Devi, Scott N. Mueller

## Abstract

With the ever-increasing complexity of microscopy modalities, it is imperative to have computational workflows that allow researchers to process and perform in-depth quantitative analysis of the resulting images. However, workflows that enable flexible, interactive and intuitive analysis from raw images to analysed data are lacking. We present Cecelia, a toolbox that integrates various open-source packages into a coherent data management suite to make quantitative image analysis accessible for non-specialists.

## Main

Choosing appropriate workflows for image analysis is complex^1^. Currently, both commercial (e.g. Imaris XTension) and open-source (e.g. CytoMAP^2^) image analysis pipelines offer only very limited capability to integrate Python and R analysis modules. There is a lack of a unified, user-friendly framework that handles the entire image data processing and analysis pipeline – from cell segmentation to data processing, analysis, visualisation – with a modular design that enables various open source as well as custom written packages to be incorporated into such workflow. With increasing multiplex imaging techniques, such as co-detection by indexing (CODEX)^3, 4^, there is a need to provide simple software solutions to spatially resolve cell populations across a broad range of experimental conditions. Software to analyse these data are currently limited to 2D images (open source: QuPath, commercial: TissueGnostics, HALO). One approach to analyse spatial tissue organisation in 3D is histo-cytometry^2, 5, 6^, which involves significant manual processing and several unconnected software packages. Computational frameworks allowing researchers to more easily conduct and manage unbiased data analysis are therefore required^2^.

Beyond static imaging, intravital microscopy enables insights into immune cell dynamics which cannot be captured from fixed samples^7, 8^. Analysis typically involves assessment of cell velocity and migration patterns ^9^ or contact durations^10^. These approaches miss the inherent heterogeneity within immune cell behaviours. Recent studies utilised manual gating^11^ or track clustering^12^ to uncover different immune cell behaviours. While these approaches enhance tracking analysis, they do not provide straightforward software frameworks to apply the same principles on other images.

To integrate static and live cell image analysis with modular workflows in a unified framework with a graphical user interface we developed Cecelia (*<u>Ce</u>*ll-*<u>cel</u>*l *i*nteraction *<u>a</u>*nalyser). This toolbox combines R/Shiny and the image viewer napari^13^, enabling real-time visualisation of resulting data analyses on the microscopy images (Figure 1a-c, Extended data Figure 1, Supplementary Notes 1-3). Cecelia is a centralised management system that allows reproducible and customised workflow design from raw images to data quantification (Figure 1a,b). Cecelia is available as R package (https://github.com/schienstockd/cecelia) and Docker container (https://github.com/schienstockd/ceceliaDocker) (Supplementary Note 4). Documentation and tutorials are available at https://cecelia.readthedocs.io/. A step-by-step walkthrough of the analysis for each image in this article is at https://cecelia.readthedocs.io/missions. The proposed framework accepts all Bio-Formats (https://www.openmicroscopy.org/bio-formats/) supported images and guides the user through segmentation (Supplementary Notes 5-7), population definition (Supplementary Notes 8-10), spatial analysis (Supplementary Notes 6, 11), and data plotting (Supplementary Note 12) (Figure 1d-f).

**Figure 1:**
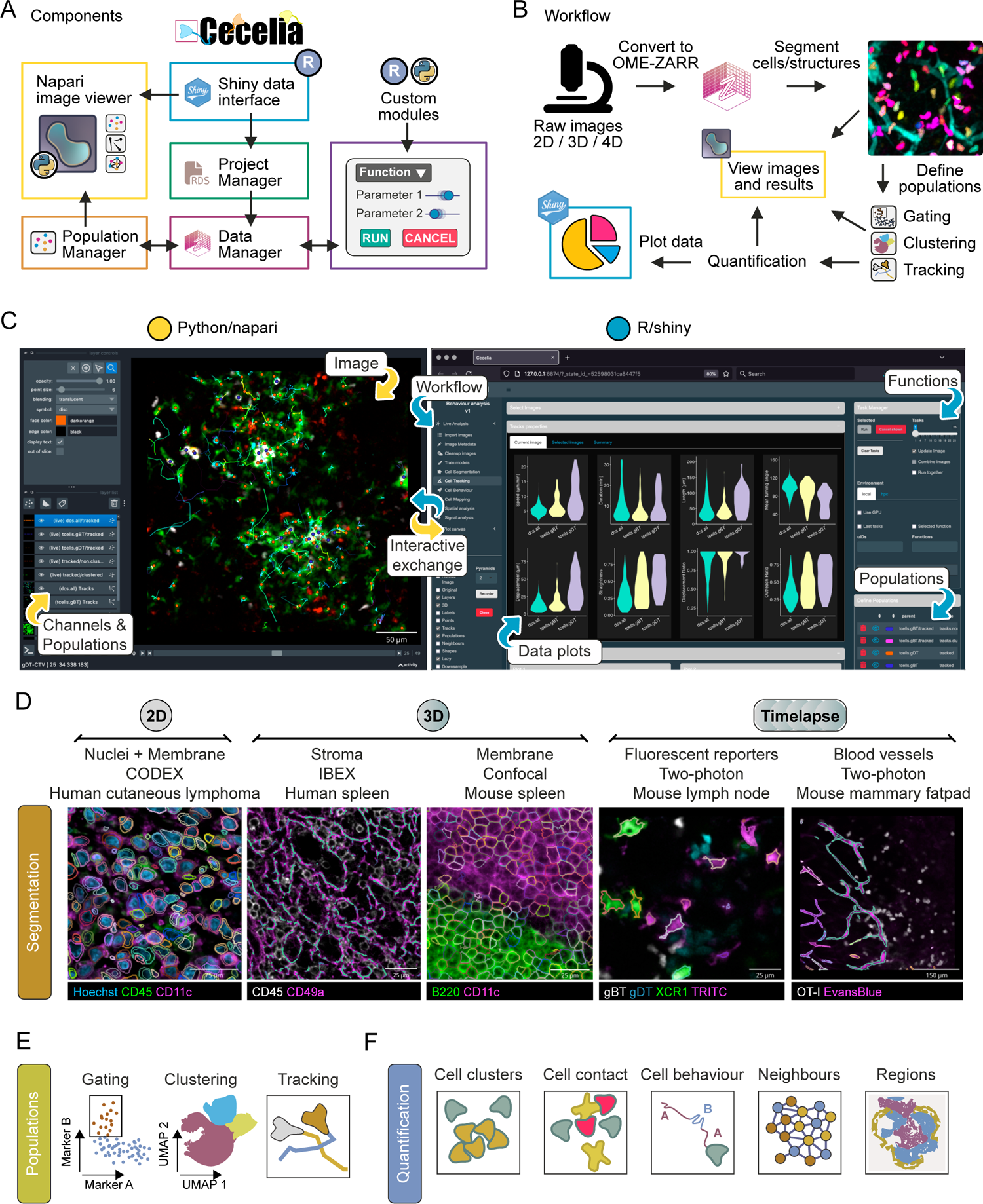
Cecelia as a general-purpose image analysis framework. a) Simplified components diagram of Cecelia’s main framework. See Extended data Figure 1b for a detailed version. b) Overview of the Cecelia workflow. Images are converted to OME-ZARR, segmented and populations defined via various means. These populations can be spatially quantified, and the results visualised on the image itself and as data plots. c) Overview of the graphical user interface. Python/napari and R/shiny are the connecting partners that form the analysis framework. d) Example images and segmentation results for various 2D, 3D and 4D imaging modalities to demonstrate the broad capability of the Cecelia framework. e) Available analysis methods available to define populations in static and live cell images. f) Main spatial analysis methods available for image quantification.

Here we applied Cecelia on a range of imaging datasets (from multidimensional high-plex static to dynamic intravital two-photon imaging). The computational packages underlying these workflows are specialised Python and R packages to demonstrate the capability of Cecelia to integrate a vast library of analysis modules. Cecelia allows independent processing of channels and cell types which can be merged into a single segmentation to process images with mixed cell sizes, shapes, markers and reporters. We applied this combined segmentation workflow to 3D spleen tissue images from mice with XCR1-venus dendritic cells and fluorescently labelled lymphocytic choriomeningitis virus (LCMV)-specific P14 T cells (Figure 2a). After segmentation, populations were manually gated (Extended data Figure 2a) and individual cell and virus interactions quantitated by Delaunay triangulation neighbour detection (Figure 2a). Our spatial analysis revealed that clustering P14 T cells preferentially interacted with XCR1^+^ DCs, whereas interactions with LCMV infected cells involved non-clustering T cells.

**Figure 2:**
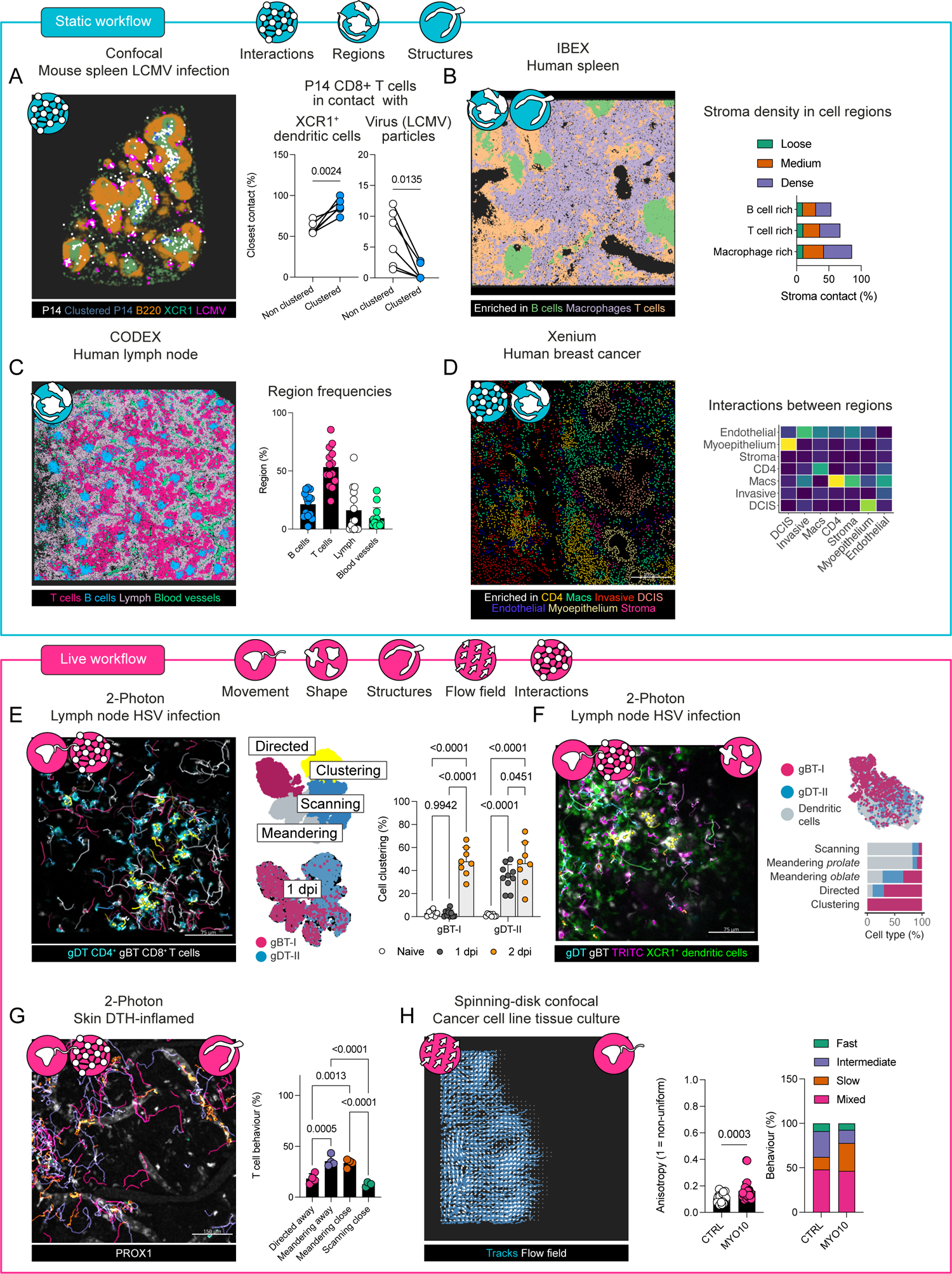
Principal workflow components of Cecelia for static and live cell image analysis a) Interactions of virus specific CD8^+^ T cells, XCR1^+^ DCs and virus infected cells in mouse spleen 1.5 days after LCMV infection (n = 6, t-test). b) Interactions of cell regions with stromal networks in healthy human spleen stained using the IBEX method. c) Cell region frequencies in healthy human node stained using CODEX (n = 14). d) Cell region interactions calculated from Xenium in situ data obtained from human breast cancer samples. e) Analysis of the behaviour of T cells in lymph nodes draining the site of skin HSV infection (n = 8-10, two-way ANOVA with Tukey’s comparison test). f) Behavioural profiling of T cells and DCs in lymph nodes 2 days after HSV infection (n = 14). g) Behaviour of T cells in DTH-inflamed skin (n = 4, two-way ANOVA with Tukey’s comparison test). h) Cancer cell migration imaged in vitro. MCF10DCIS tumour cells were treated with CTRL shRNA or MYO10 shRNA and imaged by spinning disk confocal microscopy (n = 29-30, t-test).

Next, we analysed a publicly available IBEX dataset from healthy human spleen ^14^ (Figure 2b). Cells were segmented based on membrane markers and Hoechst resulting in a combined membrane and nuclear segmentation. The segmented objects were subjected to Leiden clustering for unbiased cell population identification (Extended data Figure 2b). Cell regions were generated using KMeans, resulting in B cell, T cell and macrophage rich regions. Structural markers, including Vimentin and CD163, were used to segment the stromal network with a custom Cellpose model. To quantify where cells were residing within the stromal network, the nearest stroma branch to any one cell was determined using the k-nearest-neighbours (knn) package (k = 1) in R. The three cell regions correlated with the branching density of the stroma network with B cell regions having the least and macrophage regions the most stroma contact (Extended data Figure 2c).

To explore functional regions, we examined CODEX images from healthy human patients (portal.hubmapconsortium.org). After Leiden clustering (Extended data Figure 2d), distinct regions for T and B cells as well as lymphatics and blood vessels were identified (Figure 2c). Interactions between regions were also examined in a human breast cancer sample from Xenium in situ analysis (Figure 2d, Extended data Figure 2e). This technology allows researchers to identify RNA transcripts in fixed tissue samples. The resulting data are point-clouds of transcripts with nuclear stains such as Hoechst. This transcript data was imported as individual image channels with a pixel summation to expand the transcript signal. These examples demonstrate the capacity of the Cecelia framework to be useful for a wide variety of imaging formats.

Next, we applied our framework to dynamic 4D imaging (3D over time). Dynamic cell behaviours and interactions are not captured during static imaging but can be captured using live imaging, including intravital imaging of cells in live animal models. Basic tracking measurements are commonly used for analysis of immune cells, including cell velocity, direction and angle. We illustrate an alternative workflow using Cecelia on intravital 2-photon imaging data from the draining lymph nodes of mice during the response to cutaneous HSV infection ^15^. We imaged CD4^+^ T cells aggregating 1 day after infection, indicating antigen-specific interactions with antigen presenting cells that lead to T cell activation (Extended data Figure 3a). CD8^+^ T cells aggregated on day 2 after infection. Using Python’s btrack package to track segmented T cells, we utilised Hidden Markov Models (HMM) and Leiden clustering to extract cellular behaviour (Figure 2e). HMM are statistical models that can be used to describe sequences of observations based on underlying variables that are inherent but may not be apparent in the data. We utilised speed and angle of the individual objects to identify HMM states which were used in combination with track statistics for Leiden clustering. This method extracted considerably more refined behaviour patterns than tracking statistics alone (Extended data Figure 3b). We further demonstrate Cecelia’s utility by segregating different immune cell types within HSV infected samples, dendritic cells and T cells, by behavioural characteristics and shape differences alone (Figure 2f). This unbiased analysis of cell behaviour using Cecelia has the potential to reveal subtle differences in individual cells amongst cell populations imaged in a variety of conditions and scenarios.

Structures, such as lymphatic vessels, can influence how and where cells migrate in tissues. We analysed movies from a publicly available dataset utilising intravital 2-photon imaging of the mouse ear during Delayed-Type Hypersensitivity (DTH) response by CD4^+^ T cells to ovalbumin^16^ (https://www.immunemap.org/). Incorporating the distance of T cells to lymphatics revealed distinct modes of behaviour close to or away from these structures – with directed motion further away from and scanning closer to lymphatics (Figure 2g, Extended data Figure 3c).

Individual track measurements are informative yet cannot capture collective cellular motion. We utilised publicly available *in vitro* migration assays from a breast cancer cell line, MCF10DCIS, that either expressed a CTRL shRNA or a shRNA targeting the filopodial protein MYO10 ^17^ (Extended data Figure 3d). We extracted a flow field of cell tracks using ILEE ^18^. These flows were assessed for anisotropy, non-uniform distribution, which was increased in MYO10 targeted samples indicating increased collective directed motion while track clustering further indicated decreased migration velocity (Figure 2h). This combination of track clustering, structures and collective behaviour highlights diverse migration patterns in various imaging modalities.

We therefore demonstrate that Cecelia is a powerful and versatile framework for general image analysis based on widely used software packages. Importantly, a graphical user interface and integration of the image view napari provide a user-friendly platform for a wide range of users. Further, it is possible for users with basic programming experience to expand processing and analysis modules in the framework by adding additional packages, and for more advanced users to work directly within R or Python notebooks. We anticipate that this toolbox will facilitate use of more comprehensive image analysis approaches by a broad range of biological researchers who are non-specialists in image analysis.

## METHODS

### Mice and Infections

C57BL/6, gBT-I ^19^, gBT-I.uGFP, OT-I ^20^, OT-I.uGFP, P14 ^21^, P14.ubTomato and XCR1-venus ^22^ mice were bred in the Department of Microbiology and Immunology, The University of Melbourne. gBT-I mice encode transgenes expressing a T cell receptor recognizing the HSV-1 glycoprotein B-derived epitope gB498-505. OT-I mice encode transgenes expressing a T cell receptor recognizing OVA257-264 peptide. P14 mice encode transgenes expressing a T cell receptor recognizing the LCMV glycoprotein33-41 peptide. Animal experiments were approved by The University of Melbourne Animal Ethics Committee.

### Flank scarification, HSV infection and TRITC dye painting

Mice were anesthetised with 1:1 ratio of Ketamine (Parnell Laboratories) and Xylazil (Troy Laboratories) (10 μg per g of body weight) via intraperitoneal injection. Moisturising eye gel (Alcon) was applied to the eyes to prevent dehydration. Mouse were shaven that the left flank and belly region were exposed. Veet cream (Reckitt Benckiser) was applied using a cotton bud to aid hair removal. Veet was removed using a wet tissue and dried using a dry tissue. Location for skin abrasion was marked with a marker pen just above the tip of the spleen on the lower left flank region. For experiments marking skin migrating cells, Tetramethylrhodamine (TRITC; Life Technologies) was dissolved in 2 μl Dimethyl sulphoxide (DMSO; Sigma Aldrich) and subsequently diluted in 200 μl acetone (Chem Supply). 10 μl of this mixture was applied between the marked abrasion area and the inguinal LN and allowed to dry for about 5 min.

Skin was lightly abrased by using a Dremel for 12 sec to remove the epidermis. 10 μl of 10^6^ pfu HSV was applied to the abrased skin region. A strip of the adhesive Opsite Flexigrid (Smith+Nephew) was attached onto the infected area to contain the virus infection. Mice were bandaged with Micropore (3M) and Transpore (3M) and kept was on a heating pad for about 2 h until recovered. Bandages were removed 2 days after infection.

### LCMV infections

LCMV virus was reconstituted at 2 x 10^5^ pfu per 200 μl which were injected into mice intraperitoneally.

### T cell enrichment, labelling and adoptive transfer

To enrich naïve T cells, LNs (inguinal, popliteal, brachial, axial, mesenteric and liver) and spleen were harvested into phosphate buffered saline (PBS) with 2.5% Fetal Calf Serum (Gibco) (PBS-2.5) on ice. Organs were mechanically disrupted by using a plunger from a 1 ml syringe as a mortar on a 70 μm cell strainer. Disrupted organs were topped up with PBS-2.5 and washed. For spleens, red blood cells were lysed with 1 ml red cell lysing buffer for 5 min and washed in 10 ml PBS-2.5. For further processing, spleen and LN suspensions were combined.

To enrich CD8^+^ gBT-I, OT-I or P14 T cells, the cell suspension was incubated on ice for 30 min in a CD8^+^ negative selection antibody mix (anti-erythrocytes [Ter119], anti-I-A/E [M5/114], anti-CD4 [GK1.5], anti-Gr1 [RB6-8C5], anti-Mac-1 [M1/70], anti-F4/80 [F4/80]).

After incubation, cell suspensions were washed and resuspended in 6 ml (washed) BioMag Goat Anti-Rat IgG (Qiagen) for 20 min at 4 °C on an angled rotor. After incubation, CD8^+^ cells were enriched by running the cell suspension through a magnetic column. Enrichment purity was determined by flow cytometry with CD8α and Vα2 which typically ranged over 90%.

### Dye labelling and adoptive transfer

To label cells with CellTrace violet (CTV; Life Technologies) (5 μM stock), cells were resuspended at 10^6^/ml in PBS with 0.1% bovine serum albumin (BSA; Sigma Aldrich). 1 μl of CTV was added per 10^7^ cells for up to 10 min in a 37 °C water bath. After incubation, stained cells were washed in PBS.

To label cells with CellTracker deep red (CTDR; Life Technologies) (1 mM stock), cells were resuspended at 10^6^/ml in PBS. CTDR was prepared 1:1 with diluted pluronic (Thermo Scientific; 1:10 in PBS). 1 μl of this prepared solution was added per 10^7^ cells and incubated for 15 min in a 37 °C water bath. After the first incubation, 10 ml PBS were added for a second round of incubation for 15 min in a 37 °C water bath. At the end of this last incubation, cells were washed in PBS. Before adoptive transfer, viable cell number was determined by trypan blue staining on a haemocytometer. 0.5 – 1.0^6^ T cells were typically transferred intravenously in PBS.

### Confocal microscopy

Spleens were harvested directly into 2% Paraformaldehyde (PFA; ProSciTech) and fixed overnight at 4 °C on an angled rotor. The next day, fixing solution was removed by washing the spleens on a shaker in PBS for 5 min. After washing, spleens were transferred into a 30% sucrose solution and transferred to 4 °C for 24 h. After the tissues settled to the bottom of the tube, spleens were cut into three pieces (cross-sections) and frozen in OCT compound (Trajan Scientific) using liquid nitrogen and kept at −20 °C until needed.

Cryosections were cut at 25 μm on cryostat and air dried in dark for at least 10 min. Dried sections were fixed in acetone for 5 min and air dried for 10 min. Each cross-section was marked with a square using a hydrophobic barrier pen before applying a drop of protein block for 10 min. Antibody mixes were prepared in 2.5% Normal Donkey Serum (NDS; Sigma-Aldrich) and left on ice in dark until needed. After protein block incubation, sections were cleaned from protein block solution (Agilent Technologies) and transferred to incubation chamber (Tupperware box with foil). 50 μl antibody solution was applied to each section and samples incubated overnight at 4 °C. Antibodies used: B220 (Pacific Blue, RA3-6B2, BioLegend), CD3e (AF 700, eBio500A2, Thermo Fischer Scientific), LCMV NP (AF 594, VL-4, Bio X Cell) and CD11b (BV 421, M1/70, BioLegend).

The next day, samples were washed twice in PBS for 5 min and mounted using ProLong Gold Antifade (Thermo Fisher Scientific). For spectral imaging, single colour controls were prepared, and images acquired in lambda mode as per manufacturer instruction (Zeiss LSM 780) with a lambda bin width of 17.8 nm. Images were unmixed using ZEN blue (Zeiss, version 2.1) and further processed using Cecelia.

### Intravital two-photon microscopy

Inguinal LNs were imaged as described elsewhere ^15^. In brief, mice were anaesthetised with isoflurane (Novachem; 2.5% to start and 1.0-1.5% to maintain and supplied with an 80:20 mixture of oxygen and air) and prepared for intravital imaging. If mice were not already shaved from a previous HSV flank infection, their shaved on their belly and left hind flank and remaining hair was removed using Veet. The first incision was made just right from the midline from the bottom of the belly to top of the ribcage through the dermis. The second incision was made following the vein of the left leg to the foot and further to the base of the tail. To disconnect the connective tissue, surgical scissors were used as weights while cutting – resulting in a ‘tissue flap’.

Vetbond tissue adhesive (3M) was used to glue the ‘tissue flap’ dermis side down onto a stainless-steel platform. The inguinal LN was exposed by cutting away skin as a square window which was flooded with PBS to prevent dehydration.

Fat tissue was removed using micro scissors and gauze was utilised to remove eventual bleeding caused by small vessels within the fat tissue. Once the LN was cleared, a border of grease was applied to the square cut-out which could hold a pool of PBS. A square coverslip (ProSciTech) was placed on top, and a hydrophobic barrier pen (Sigma Aldrich) was used to draw a square on top of the coverslip where water could be placed for the water objective of the two-photon microscope.

Mice were placed in a custom build heated (35 °C) humidified chamber for two-photon imaging with an Olympus FV-MPERS multiphoton system and a 25x water objective. The system was equipped with two lasers Mai-Tai and Insight DeepSee which were used at 820 nm (for CTV) and 980 nm (for GCAMP, CTDR and or TRITC) wavelengths respectively. The time interval between frames was 10 sec, with a 5 μm gap between optical planes. Images were processed using Cecelia.

### Image analysis

An overview of used packages is listed in Supplementary table 1. For tutorials and user interface interaction, the reader is referred to https://cecelia.readthedocs.io. Images were imported as OME-Zarr ^23^ using bioformats2raw (Glencoe Software). For Xenium files, point data was converted into images with a local sum of pixels to aggregate and make transcripts easier to view and analyse.

Images underwent ‘cleanup’ routines if needed to account for overspill between individual channels. This is especially important for two-photon images as they contain, generally, a significant amount of autofluorescence and channels spill-over. While correction in Imaris is often done using channel subtraction, we opted for channel division instead as this resulted in cleaner output. Subsequently images underwent cell and structure segmentation. While we integrated various deep learning approaches into the framework, Cellpose ^24^ has proven to be the most practical as it is widely applicable to various cases and custom models can be generated relatively simply. Depending on whether the image is static or live the identified labels undergo population definition or tracking respectively.

Populations were defined by flow cytometry gating, implemented using flowWorkspace, or by population clustering, using Leiden clustering. For tracking, we utilised the Bayesian tracker btrack as it provided convenient integration into Cecelia. We currently have no option to manually correct the resulting tracks. Tracking statistics were computed with celltrackR. Populations and tracked cells were further utilised for spatial analysis.

As the number of cells in static images can be rather large, most analysis was done based on centroids with packages such as DBSCAN. For two-photon analysis, 3D meshes were generated using trimesh which also provides a collision manager to detect cell interactions. For dynamic cell behaviour, the HMM functionality from depmixS4 was utilised and combined with Leiden clustering. Data plots were generated in R or GraphPad Prism 10 (GraphPad Software).

## Data availability

Images not generated in this article were downloaded from the sources outlined in the Supplementary table 2.

## Code availability

Code can be accessed at https://github.com/schienstockd/cecelia.

## Acknowledgements

This research was supported by The University of Melbourne’s Research Computing Services and the Petascale Campus Initiative. This work was supported by a National Health and Medical Research Council Investigator Fellowship to S.N.M. (2017220). We thank the Biological Optical Microscopy Platform at the University of Melbourne for access to microscope facilities and the Biological Research Facility for management, breeding and maintenance of mice.

## Author contributions

D.S. and S.N.M. conceived the project. D.S. performed experiments, data analysis, computational implementation and software development. J.L.H. and S.D. provided experimental data and provided feedback on the manuscript. D.S. prepared the manuscript and D.S. and S.N.M. edited the manuscript. S.N.M. supervised the entire project.

---

**Extended data Figure 1:**
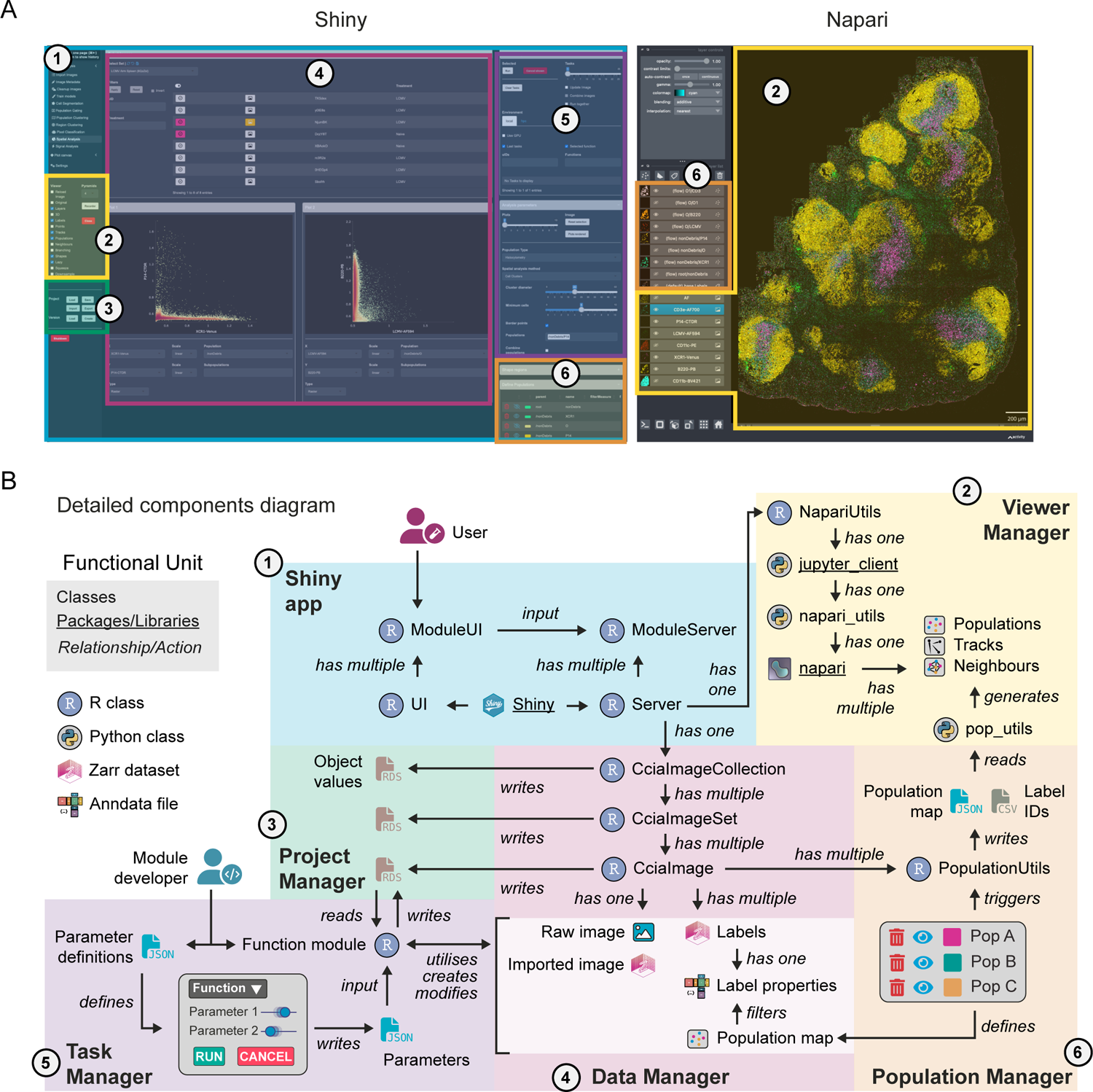
Main components of Cecelia User interface (a) and the respective underlying software components (b). (1) The user interacts with the shiny app which is a modular construct of UI and Server components. The main shiny server has one connection to control napari (via NapariUtils) and access data (via CciaImageCollection). (2) Napari is controlled via a jupyter client and a custom napari utilities module. In this way populations, tracks, neighbours, etc. can be toggled from R via Python. (3) The project management class triggers read and write processes of the individual image analysis and the main project values. (4) Each image record has the original raw image file as well as modified versions that the user generates. Each image also contains quantification and further analysis executed by the user. (5) The module developer can provide functions in R and Python to server user requirements. The parameters for each function module can be exposed to user input via structured JSON files. (6) Populations can be generated via different means. The main route is via the PopulationUtils class which holds the population information and can write out structured JSON files and labels identifiers that can be read and displayed by napari.

**Extended data Figure 2:**
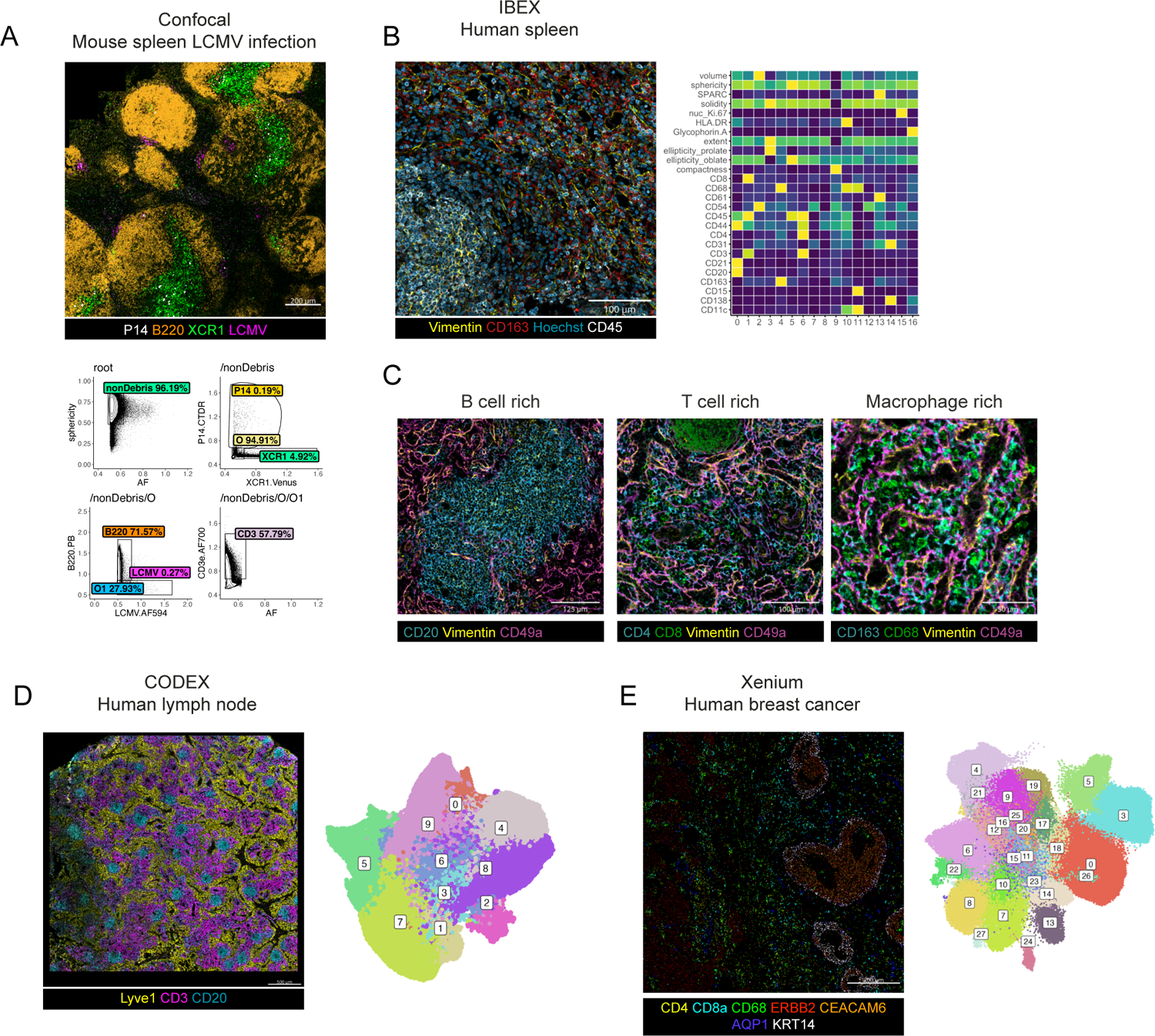
Examples of static image analysis capabilities of Cecelia a) Processing and gating of mouse spleen samples after LCMV infection. Other populations: O = Others; O1 = Others 1. b) Processing and cluster extraction from human spleen samples. c) Examples of cell regions with their respective stromal cell networks. d) Processing and cluster extraction for human lymph node. e) Processing and cluster extraction for human breast cancer samples.

**Extended data Figure 3:**
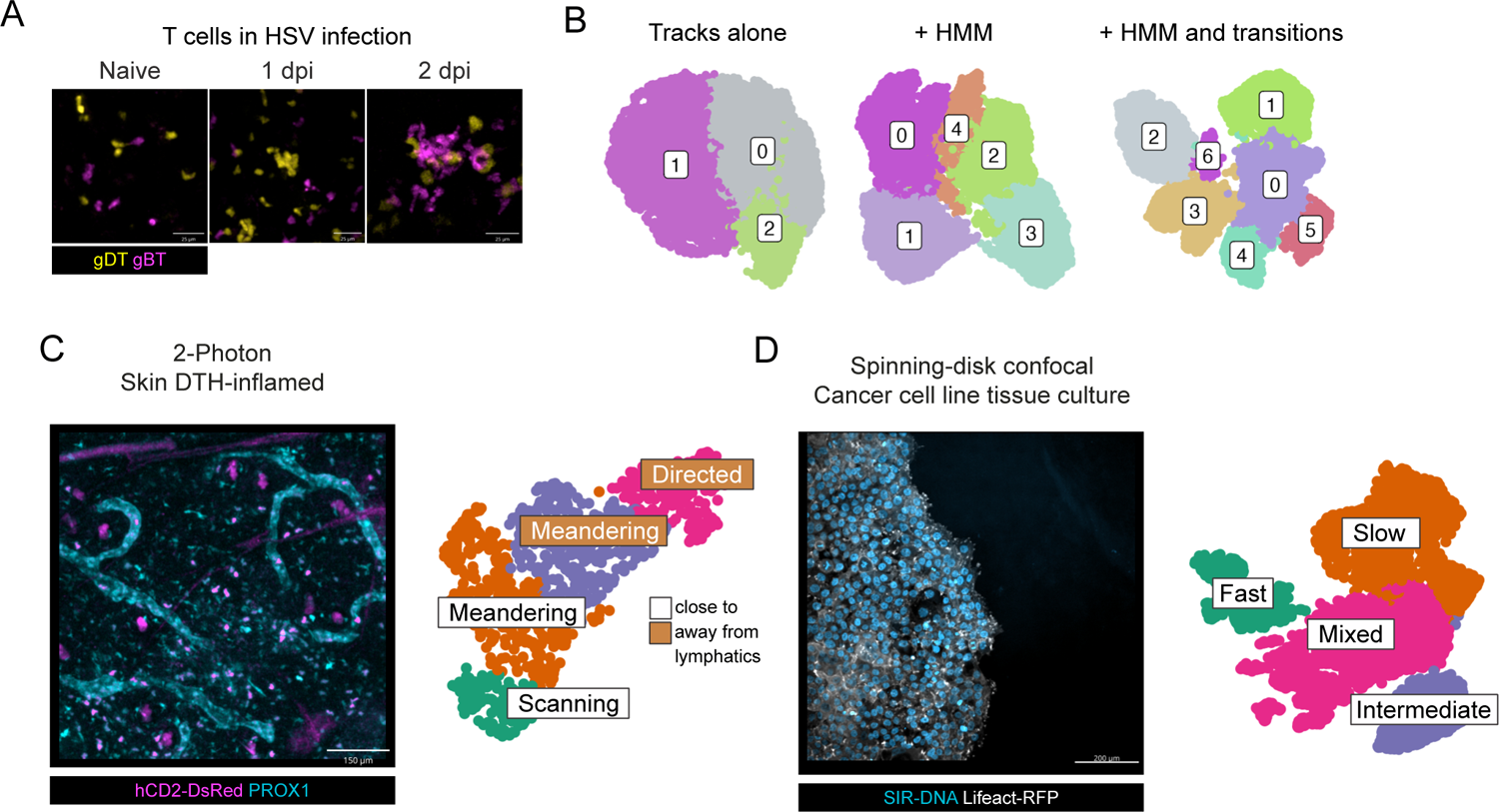
Live cell image analysis and behavioural cell profiling using Cecelia a) Example images of T cell clustering in the lymph node after HSV infection imaged using 2-photon microscopy. b) Resulting cell behaviour clusters with either tracks alone or with inclusion of hidden Markov models (HMM) or with HMM and respective state transitions. c) Processing and cluster extraction for T cell behaviour in DTH-inflamed skin samples imaged using 2-photon microscopy. d) Processing and cluster extraction for cancer cells after knockdown of the filopodial protein MYO10 imaged using spinning disk confocal microscopy.

