## Supplementary Notes for "Cecelia: a multifunctional image analysis toolbox for decoding spatial cellular interactions and behaviour"

**Supplementary Table 1: Main packages used in the Cecelia framework**

| Usage | Package | Language |
|---|---|---|
| Data storage | anndata | Python |
| Tracking of labels | btrack | Python |
| Cell segmentation and image denoising | cellpose | Python |
| Tracking measurements | celltrackR | R |
| Concavity for cell areas | concaveman | R |
| Viewing images in napari | dask | Python |
| Cell clusters and contact | dbscan | R |
| Hidden Markov Models | depmixS4 | R |
| Gating of cell populations | flowWorkspace | R |
| Bridge between R and Python | jupyter | Python |
| Population clustering | leidenalg | Python |
| Image denoising | n2v | Python |
| Viewing images | napari | Python |
| Image metadata | ome-types | Python |
| Managing processes | parallel | R |
| Interactive plotting and cell population gating | plotly | R |
| Population clustering | scanpy | Python |
| General image processing | scikit-image | Python |
| Data management and processing | Shiny | R |
| Network branch extraction | skan | Python |
| Spatial statistics, for example population distributions | spatstat | R |
| Neighbour analysis | squidpy | Python |
| General working with image loading/saving | tifffile | Python |
| 3D mesh generation, shapes and contact detection | trimesh | Python |
| Imaging data | zarr | Python |

**Supplementary Table 2: Datasets used for figures**

| Figure | Description | Reference |
|---|---|---|
| 1b | Confocal mouse spleen | This paper |
| | CODEX Human cutaneous lymphoma | [1] |
| | Two-photon Mouse lymph node | [2] |
| | Two-photon Mouse mammary fatpad | This paper |
| | IBEX Human spleen | zenodo.org/records/4632320 |
| 2a | Confocal Mouse spleen LCMV infection | This paper |
| 2b | IBEX Human spleen | zenodo.org/records/4632320 |
| 2c | CODEX Human lymph node | portal.hubmapconsortium.org |
| 2d | Xenium Human breast cancer | 10xgenomics.com/products/xenium-in-situ/human-breast-dataset-explorer |
| 2e | Two-photon Lymph node HSV infection | [2] |
| 2f | Two-photon Lymph node HSV infection | This paper |
| 2g | Two-photon Skin DTH-inflamed | app.immunemap.org/experiment-public-view?id=49 |
| 2h | Spinning-disk confocal Cancer cell line tissue culture | zenodo.org/records/10539020 |

**Supplementary Note 1 – Rationale for implementation of the approach**

Image analysis is often a combination and integration of different tools, packages and frameworks from various programming and scripting languages. This diversity is good to increase the efficiency and ease of use for specific scenarios and user groups. The downside of this approach is a proliferation of various tools and packages that researchers must traverse to analyse their images. We therefore aimed to develop a toolkit that would support research questions specific to the immunological research in our lab but would also support cell biology research in general.

Python has been adopted by many image analysists due to its programming accessibility, vast array of processing libraries and relative ease for non-programmers to obtain reasonable automation of results. The recent development of a multidimensional image viewer in python, termed napari [3], has expanded this accessibility to image processing and opened up the possibility to implement custom workflows relatively fast within a simple user interface. Within napari, users have the capacity to develop custom plugins for specific or more generic applications which resulted in numerous custom plugins in a relatively short time (https://www.napari-hub.org/). With the advent of the multi-core and parallel computing framework Dask (https://www.dask.org/) and the Next Generation File Format (NGFF) from the Open Microscopy Environment (OME) [4], new and more accessible options are now available to parallelise processing steps and to store image information in an open and extensible community supported image file format. Downstream of image processing, respective data analysis is dependent on expertise in the research group. Popular choices include python, MATLAB and R. Python and R have similar analysis capabilities and are in many cases interchangeable.

The question in this context is, whether and how these tools could be integrated into a generic toolbox for cell biologists to conduct image processing and downstream analysis within the same workflow. We opted for napari as the main image viewer because of its increasing popularity and sustained support by the Chan-Zuckerberg Initiative (https://chanzuckerberg.com/science/programs-resources/imaging/). The downside of napari is that it must run locally. With the advent of more web-frameworks and remote cloud computing, there are a variety of web-frameworks available to view and analyse images [5, 6]; however, we could not find a suitable image viewer that runs in a web-environment and supports 2D, 3D and 3D+time images with the option to display images larger than commonly available local memory and cell tracks.

Napari alone however might not be suitable for an extensive framework to process, manage and analyse imaging data as its primary use is viewing images. We therefore considered several options to combine image processing and viewing using napari with the ability to interactively generate data plots and images together with downstream analysis – these include: a native napari plugin with plotting engine, napari embedded into another python Qt-based GUI and napari in combination with a reactive web-framework. The option to add use case specific plugins to napari is very powerful. Images can be directly linked to simple plots, such as clustering segmented objects and viewing them interactively (https://github.com/BiAPoL/napari-clusters-plotter). While these approaches will elevate how researchers can work and interact with images, our aim was to provide a comprehensive toolbox that supports different kinds of plots and tables in an interactive manner with the possibility to extend this as different use-cases arise in the future without the need to reprogram visual aids but rather to make use of the variety of packages available in any scripting or programming language.

As napari is a native Qt application (a cross-platform framework to create user interfaces, https://www.qt.io/), it could also be embedded into a another Qt-based application as done for the application PartSeg which was designed to segment structures within nuclei [7]. This solution would add more interactivity around napari but would also require the implementation of a significant amount of interactive plotting functionality which is currently available within Matplotlib (https://matplotlib.org), albeit limited. Reactive web frameworks such as Dash (Python) and Shiny (R) have significantly reduced the barrier to display data interactively with a reduced amount of implementation work required. Dash and Shiny have similar functionalities and utilise the same plotting engines such as Plotly (https://plotly.com/). Shiny's reactive programming capability can reduce the amount of code required to produce the same output while also enabling to store intermediate variables which is not possible in Dash. R further has the advantage to have a strong bias towards statistical analysis packages and tests. Keeping data analysis and image viewing in separate 'compartments' therefore opens up the space to benefit from both approaches, that is, napari's vast plugin system and the wide variety of interactive libraries available for reactive web-frameworks.

**Supplementary Note 2 - Design and Implementation**

Cecelia consists of a user interface in R/Shiny and python/napari (Supplementary Figure 1a). The main user interface for image, task and population management was implemented in R while image processing, segmentation and other related functions were implemented in python. The main components are structured into 1) module pages, 2) napari viewer managements, 3) project management, 4) image management, 5) task management and 6) population management (Supplementary Figure 1b). The user mainly interacts with the module pages from Shiny while the individual tasks can be provided by custom modules which are implemented in R and/or python depending on the functionality needed.

A Jupyter kernel serves as the communication platform between Shiny and napari. The way back from python to R is utilising JSON files that are written by napari and read into Shiny using a *reactivePoll*, for example when the user selects points on the image during gating to show them on the gating plot. For image processing, we utilised the recently developed Open Microscopy Environment Next Generation File Format (OME-NGFF) [4] which is based on small compressed files in combination with Dask for lazy loading and processing of large scale images. Every image is represented as a reactive persistent object (RPO) within R. 'Reactive' means that the object acts as any other reactive variable within Shiny. This design was based on a theoretical description within the R/Shiny community (https://community.rstudio.com/t/good-way-to-create-a-reactive-aware-r6-class/84890/8). 'Persistent' means that the information of every object is saved to individual RDS files. Every object is therefore saved independently and can be moved between computing environments to process data in different physical locations. For all processing steps, especially deep learning (DL) based segmentation, the user has the option to run every operation on a High-Performance Computing (HPC) system with the advantage of increasing computation, memory and graphics card resources.

For each unique RPO, there is a corresponding folder with its unique identifier to store all object related data and analysis. Every project, which also has a unique identifier, holds corresponding experimental sets with individual images. A project can contain a series of experiments that are related to each other and require similar sets of analysis workflows. Projects are stored at the root of the folder structure. Each project has several versions, where version '0' contains all image information and versions '1' to 'n' contains all the processed information. Versions can be created by the user as a backup to test whether the experimental data could be analysed differently by saving a specific state of the already analysed data. Within each data folder, there are several subfolders that contain segmentation labels and properties as well the population definitions.

**Supplementary Note 3 – Task management**

Each RPO can be used to process tasks. A major drawback of using R for process management is that there is no support for concurrent threads. Individual tasks can however run in separate parallel processes. Within Shiny, there is currently no direct module or library that manages processes. Our aim was to provide a framework that could utilise R and python packages within tasks that can run locally as well as remotely on HPC systems. The current implementation works with the R package mcparallel to run individual forked processes; an approach which only works on Unix systems.

The current implementation is based on two main classes: *taskLauncher* and *taskProcess*. *taskLauncher* is a task container which provides functions to start, stop and retrieve results. All tasks inherit from *taskProcess* which provides functions to retrieve the given parameters and work with the RPO of the current task. Generally, each task is coupled to an RPO where logfiles and results will be written to. When multiple images are being processed, the experimental set of these images will be taken as the processing RPO. For tasks running on the HPC, the current RPO state file (RDS file) will be uploaded together with the task parameters. The HPC job will start an R script to run the task.

Processing within python is done by calling corresponding python scripts in R via system calls. Certain features, especially Dask, are currently not working when using the R package reticulate which is used to run python libraries within R, hence python scripts are called directly rather than integrated within R. Python tasks are handled like R tasks in the sense that parameters are passed to the script via JSON files.

Every task function has a unique name across the app; for example, Cellpose segmentation is called *segment.cellpose*. This name is used to link functions to individual module pages within Shiny; that means, all tasks starting with *segment* will be shown on the segmentation module page. Processing parameters for every function are defined in JSON files. Based on the input definitions within these JSON files, the input parameters to be shown in the Shiny app for every function are generated and then passed to the corresponding function call. Parameters passed in this way are then available in every process by calling the function *self$funParams()*. In this way, it is relatively simple to introduce and implement new functionality within the app. The developer must define a JSON file for parameters and an R file for the function implementation. Commonly used functionality such as adding populations, running tasks or plotting data in flow cytometry plots is also implemented in this modular way. The app is therefore extendible beyond the current available modules.

**Supplementary Note 4 – Docker setup**

The app utilises a wide range of packages, libraries and several external programs. We have aimed to make the installation as minimal as possible by providing an R package structure with integrated python scripts. The user then must run a series of initial methods to install further package dependencies and retrieve DL models. For ease of usage, we provided a contained running environment using a Docker container. It is currently not easy to get napari working within a Docker environment. We have therefore chosen to install napari within a conda environment on the host. This environment communicates with Shiny which is packaged within the Docker container. This setup is currently required for Windows users.

**Supplementary Note 5 – Image segmentation**

DL based methods are arguably the most efficient and effective means for image segmentation [8-10]. Whole-slide 2D images with nuclei and membrane staining can now be relatively reliably segmented and analysed even in crowded conditions. Our ambition was to provide workflows that could also segment larger images in 3D and 3D+time with varying cell sizes and shapes without the user having to retrain models. We have therefore incorporated the commonly used DL frameworks Mesmer [8], Stardist [9] and Cellpose [10]. For our purposes, Cellpose has been the most useful as it can segment a variety of cell shapes and sizes in 2D and 3D without the need to create a series of custom models.

**Supplementary Note 6 – Object measurements**

After cell segmentation, objects must be measured for further analysis. Mesmer [8] and a recent publication on two-photon image analysis [11] provided a wide variety of measurements for 2D and 3D object analysis respectively. These 3D measurements required the generation of surface meshes. There are several libraries available within python to create and manage meshes, we opted for trimesh (https://trimsh.org/). Trimesh can extract meshes from binary images via scikit-image's marching cube algorithm. These meshes can be saved for later use to detect object interactions which Trimesh supports via a Collision Manager. We have currently not implemented a way of combining meshes into single object files which results in every mesh being saved as a single STL file (a file format to store vertices) which takes up a significant amount of storage space for larger images and results in slow processing as meshes must be loaded individually from disk.

**Supplementary Note 7 – Object properties storage and access**

There are several options available to store data. We are using R and python together which required an approach to read and write data from both sides. We opted for the HDF5 based file-format Anndata (https://github.com/scverse/anndata) which was originally developed for single cell analysis within the python package Scanpy [12]. The data can be accessed from R via the reticulate library. For each segmentation, object measurements and properties are saved within a single Anndata file. For easier access, we implemented a helper class called *LabelPropsView* which provides functions for the most used access options. One challenge is that R and python do not operate in the same process and therefore access to these datafiles can lead to a 'race condition' where both environments might access the same file at the same time. Operations between these two systems therefore must be executed coordinated.

**Supplementary Note 8 – Population management**

We aimed to support different kinds of imaging data including tracked, gated and clustered cell populations. As each of these have slightly different requirements, we decided to split these into different classes with one common base class. Currently, all populations are defined and saved within R and accessible in python as read-only. This decision was made initially as the gated cell population class is based on the R package flowWorkspace which provides a population gating framework. The populations IDs for every cell population are written out as individual CSV files that python can read out and filter the main dataset accordingly. Other sub-populations from tracking and clustering are implemented as 'filtered' populations; for example, tracked cells have a *track_id* above 0 which is used to define the 'tracked' cell population. These filters are again applied within R which is then saving the population IDs to CSV files for python to display. Clustered populations and regions work in the same manner.

**Supplementary Note 9 – Flow gating**

One of the main functionalities that were required for the app was to be able to gate on cell populations and check the gating result at the same time on the image. There are commercial flow cytometry platforms based on web-technology, such as OMIQ (https://www.omiq.ai/) which developed their own plotting and gating engine. Our approach was to try to utilise existing components from Shiny and R to create a gating interface. Plotly (https://plotly.com/) is one of the most popular frameworks for interactive graphing and plotting which is integrated into Shiny and python's dash (https://plotly.com/dash/). Plotly enables drawing of shapes onto the data which can be utilised for creating population gates. Every time the user creates a shape, a *relayout* event is triggered by the plot. This *relayout* event contains the points data from the drawn shape which can be used to create or change the population gating.

Gating of cell populations is essentially a simple 2D shape which is drawn on top of datapoints. flowWorkspace is an R package that was designed to represent gating hierarchies and import and export data from and to the commercial and commonly used FlowJo software (https://www.flowjo.com/). The combination of these two packages, plotly and flowWorkspace, together with Shiny's reactive framework enables interactive population gating. The current implementation is however comparatively inefficient due to the nature of reactivity. Future developments should more carefully consider the dependencies and interactions of reactive variables and respective functions and whether plotly could be replaced by a custom designed plotting library for flow cytometry, as done in OMIQ, to increase efficiency and user experience.

One issue for displaying segmentation data is the large number of points with larger images. A commonly used method is to use raster images instead of plotting individual points. We utilised the R package rasterly (https://github.com/plotly/rasterly) to generate raster plots and contour plots to achieve this.

**Supplementary Note 10 – Object tracking**

Object tracking has been widely simplified with the advent of the ImageJ plugin Trackmate [13] which was recently expanded to also track results from DL segmentation frameworks [14]. Here we utilised the bayesian tracking package btrack within python [15] to track results from segmentations.

**Supplementary Note 11 – Neighbour detection**

Neighbour detection can be done in several ways, such as a fixed scanning window across the image, cell centred radii or Delaunay triangulation. Squidpy [16] was recently developed to analyse spatial single cell data and is based on Anndata. We can therefore utilise the functions therein to obtain cell neighbours using the mentioned methods.

**Supplementary Note 12 – Plotting data**

One advantage of R/Shiny is that generating data plots is relatively straightforward. Shiny will render any plot that can be produced within R as image files that are then displayed on the web-application. There is therefore a slight delay between generating a plot, saving it as an image and showing the image in the web-application. This procedure is fine for plots that do not require interactivity. For interactive plots where users can select points and drag shapes, as for population gating, the graphing library Plotly is more appropriate.

Processing and analysing imaging data has several manual and comparatively subjective steps where users must choose parameters that will influence the downstream analysis and the final figure. We therefore aimed to provide a workflow where the effects of certain parameter choices can be examined interactively. Very often the final analysis of images is only loosely bound to the original image. To display data in different ways and interactions, we have adopted a series of previously defined Shiny apps (https://huygens.science.uva.nl/) and integrated them into Cecelia within the 'Plot canvas' section. The data for the plots is taken from the analysed images which results in direct visualisation of the processed images so that users can see the impact of different processing parameters on the image as well as on the resulting data plot.
